## Supplementary material for "A workflow reproducibility scale for automatic validation of biological interpretation results": LaTeX Source: example_file_entity.pdf

```json
{
  "@context": ["https://w3id.org/ro/crate/1.1/context", { ... }],
  "@graph": [
    ...,
    {
      "@id": "outputs/star_salmon/RAP1_UNINDUCED_REP2.markdup.sorted.bam",
      "@type": ["File", "FormalParameter", "OutputFile"],
      "contentSize": 3279083,
      "dateModified": "2022-09-08T08:52:19.755363",
      "encodingFormat": "application/gzip",
      "format": {
        "@id": "http://edamontology.org/format_2572"
      },
      "gid": 1000,
      "mode": "-rw-r--r--",
      "sha512": "2d6c8436dd1da0e4e49f9bdfbf8d656d7740f7eae149bb2add417d6739c05aeb441aa84239041f9aac87688042c7b31bfca5c95d0dfaf742512f2e740a788979",
      "stats": {
        "@id": "#31d3ba80-21df-4ab7-93e7-558154d07161"
      },
      "uid": 1000,
      "url": "http://localhost:1122/runs/93f4d8bf-424d-4d5e-bc79-9482e1620be9/data/outputs/star_salmon/RAP1_UNINDUCED_REP2.markdup.sorted.bam"
    },
    {
      "@id": "#31d3ba80-21df-4ab7-93e7-558154d07161",
      "@type": ["FileStats"],
      "duplicateRate": 0.8027008887713803,
      "duplicateReads": 78936,
      "generatedBy": {
        "@id": "#samtools"
      },
      "mappedRate": 1.0,
      "mappedReads": 98338,
      "totalReads": 98338,
      "unmappedRate": 0.0,
      "unmappedReads": 0
    },
    ...
  ]
}
```
