## Supplementary material for "A workflow reproducibility scale for automatic validation of biological interpretation results": LaTeX Source: tonkaz_cli_example.pdf

```
...

=== Level3 ⭐⭐⭐ (Same Checksum, 13/25 files)

  - star_salmon/rseqc/junction_annotation/bed/RAP1_IAA_30M_REP1.junction.Interact.bed
  - star_salmon/rseqc/junction_annotation/bed/RAP1_IAA_30M_REP1.junction.bed
  - star_salmon/rseqc/junction_annotation/bed/RAP1_UNINDUCED_REP1.junction.Interact.bed
  - star_salmon/rseqc/junction_annotation/bed/RAP1_UNINDUCED_REP1.junction.bed
  - star_salmon/rseqc/junction_annotation/bed/RAP1_UNINDUCED_REP2.junction.Interact.bed
  - star_salmon/rseqc/junction_annotation/bed/RAP1_UNINDUCED_REP2.junction.bed
  - star_salmon/rseqc/junction_annotation/bed/WT_REP2.junction.Interact.bed
  - star_salmon/rseqc/junction_annotation/bed/WT_REP2.junction.bed
  - star_salmon/stringtie/RAP1_IAA_30M_REP1.coverage.gtf
  - star_salmon/stringtie/RAP1_UNINDUCED_REP1.coverage.gtf
  - star_salmon/stringtie/RAP1_UNINDUCED_REP2.coverage.gtf
  - star_salmon/stringtie/WT_REP1.coverage.gtf
  - star_salmon/stringtie/WT_REP2.coverage.gtf

=== Level2 ⭐⭐ (Similar Features, 12/25 files)

  - star_salmon/WT_REP1.markdup.sorted.bam
    .---------------------------------------------------------------------.
    |                |        in Crate1         |        in Crate2         |
    |----------------|--------------------------|--------------------------|
    | File Size      | 6.86 MB (7196616)        | 6.86 MB (7192272)        |
    | Total Reads    | 188243                   | 188241                   |
    |   # Mapped     | 188243 (100.00%)         | 188241 (100.00%)         |
    |   # Duplicate  | 38470 (20.44%)           | 38470 (20.44%)           |
    '---------------------------------------------------------------------'

  - star_salmon/rseqc/junction_annotation/bed/WT_REP1.junction.Interact.bed
    .---------------------------------------------------------------------.
    |                |        in Crate1         |        in Crate2         |
    |----------------|--------------------------|--------------------------|
    | File Size      | 22.46 KB (22995)         | 22.32 KB (22856)         |
    | Line Count     | 162                      | 161                      |
    '---------------------------------------------------------------------'

  - star_salmon/stringtie/WT_REP2.transcripts.gtf
    .---------------------------------------------------------------------.
    |                |        in Crate1         |        in Crate2         |
    |----------------|--------------------------|--------------------------|
    | File Size      | 37.82 KB (38729)         | 37.82 KB (38729)         |
    | Line Count     | 259                      | 259                      |
    '---------------------------------------------------------------------'

...

=== Level1 ⭐ (Different Features, 0/25 files)

=== Level0 (Not Found, Crate1: 0 files, Crate2: 0 files)

Summarize compare result:

  .----------------------------------------------------------------------------.
  |      Reproducibility         |   Level    |      Definition       |   File #    |
  |------------------------------|------------|-----------------------|-------------|
  | Fully Reproduced             | ⭐⭐⭐     | Same Checksum         | 13 files    |
  | Acceptable Differences       | ⭐⭐       | Similar Features      | 12 files    |
  | Unacceptable Differences     | ⭐         | Different Features    | 0 files     |
  | Not Reproduced               |            | Not Found             | 0 files     |
  '----------------------------------------------------------------------------'
```
