## Supplementary material for "A workflow reproducibility scale for automatic validation of biological interpretation results": LaTeX Source: main.bib

@article{stephens_big_2015,
	title        = {{Big data: Astronomical or genomical?}},
	author       = {Stephens, Zachary D. and Lee, Skylar Y. and Faghri, Faraz and Campbell, Roy H. and Zhai, Chengxiang and Efron, Miles J. and others},
	year         = 2015,
	journal      = {{PLOS Biology}},
	volume       = 13,
	number       = 7,
	pages        = {e1002195},
	doi          = {10.1371/journal.pbio.1002195}
}
@article{stein_case_2010,
	title        = {{The case for cloud computing in genome informatics}},
	author       = {Stein, Lincoln D.},
	year         = 2010,
	journal      = {{Genome Biology}},
	volume       = 11,
	number       = 5,
	pages        = 207,
	doi          = {10.1186/gb-2010-11-5-207}
}
@article{goodwin_coming_2016,
	title        = {{Coming of age: Ten years of next-generation sequencing technologies}},
	author       = {Goodwin, Sara and {McPherson}, John D. and {McCombie}, Richard W.},
	year         = 2016,
	journal      = {{Nature Reviews Genetics}},
	volume       = 17,
	number       = 6,
	pages        = {333--351},
	doi          = {10.1038/nrg.2016.49}
}
@article{prins_toward_2015,
	title        = {{Toward effective software solutions for big biology}},
	author       = {Prins, Pjotr and de Ligt, Joep and Tarasov, Artem and Jansen, Ritsert C. and Cuppen, Edwin and Bourne, Philip E.},
	year         = 2015,
	journal      = {{Nature Biotechnology}},
	volume       = 33,
	number       = 7,
	pages        = {686--687},
	doi          = {10.1038/nbt.3240}
}
@article{perkel_workflow_2019,
	title        = {{Workflow systems turn raw data into scientific knowledge}},
	author       = {Perkel, Jeffrey M.},
	year         = 2019,
	journal      = {{Nature}},
	volume       = 573,
	number       = 7772,
	pages        = {149--151},
	doi          = {10.1038/d41586-019-02619-z}
}
@inproceedings{banati_evaluating_2016,
	title        = {{Evaluating the reproducibility cost of the scientific workflows}},
	author       = {B{\'a}n{\'a}ti, Anna and Kacsuk, P{\'e}ter and Kozlovszky, Mikl{\'o}s},
	year         = {2016-05},
	booktitle    = {{2016 IEEE 11th International Symposium on Applied Computational Intelligence and Informatics (SACI)}},
	pages        = {187--190},
	doi          = {10.1109/saci.2016.7507367}
}
@article{noauthor_software_2014,
	title        = {{Software with impact}},
	year         = 2014,
	journal      = {{Nature Methods}},
	volume       = 11,
	number       = 3,
	pages        = {211--211},
	doi          = {10.1038/nmeth.2880}
}
@article{noauthor_rebooting_2015,
	title        = {{Rebooting review}},
	year         = 2015,
	journal      = {{Nature Biotechnology}},
	volume       = 33,
	number       = 4,
	pages        = {319--319},
	doi          = {10.1038/nbt.3202}
}
@inproceedings{drummond_replicability_2009,
	title        = {{Replicability is not reproducibility: Nor is it good science}},
	author       = {Drummond, Chris},
	year         = 2009,
	booktitle    = {{Proceedings of the Evaluation Methods for Machine Learning Workshop at the 26th ICML}},
	volume       = 1
}
@misc{acm_artifact_2021,
	title        = {{Artifact review and badging version 1.1}},
	author       = {{Association for Computing Machinery}},
	year         = 2020,
	url          = {https://www.acm.org/publications/policies/artifact-review-and-badging-current},
	urldate      = {accessed 2022-09-22},
	howpublished = {\url{https://www.acm.org/publications/policies/artifact-review-and-badging-current}}
}
@article{peng_reproducible_2011,
	title        = {{Reproducible research in computational science}},
	author       = {Peng, Roger D.},
	year         = 2011,
	journal      = {{Science}},
	volume       = 334,
	number       = 6060,
	pages        = {1226--1227},
	doi          = {10.1126/science.1213847}
}
@article{leprevost_best_2014,
	title        = {{On best practices in the development of bioinformatics software}},
	author       = {Leprevost, Felipe da Veiga and Barbosa, Valmir C. and Francisco, Eduardo L. and Perez-Riverol, Yasset and Carvalho, Paulo C.},
	year         = 2014,
	journal      = {{Frontiers in Genetics}},
	volume       = 5,
	doi          = {10.3389/fgene.2014.00199}
}
@article{wratten_reproducible_2021,
	title        = {{Reproducible, scalable, and shareable analysis pipelines with bioinformatics workflow managers}},
	author       = {Wratten, Laura and Wilm, Andreas and G\"{o}ke, Jonathan},
	year         = 2021,
	journal      = {{Nature Methods}},
	volume       = 18,
	number       = 10,
	pages        = {1161--1168},
	doi          = {10.1038/s41592-021-01254-9}
}
@misc{amstutz_existing_2021,
	title        = {{Existing workflow systems}},
	author       = {Amstutz, Peter and Mikheev, Maxim and Crusoe, Michael R. and Tijani\'{c}, Neboj\v{s}a and Lampa, Samuel},
	year         = 2021,
	url          = {https://s.apache.org/existing-workflow-systems},
	urldate      = {accessed 2022-09-22},
	howpublished = {\url{https://s.apache.org/existing-workflow-systems}}
}
@article{crusoe_methods_2021,
	title        = {{Methods included: Standardizing computational reuse and portability with the common workflow language}},
	author       = {Crusoe, Michael R. and Abeln, Sanne and Iosup, Alexandru and Amstutz, Peter and Chilton, John and Tijani\'{c}, Neboj\v{s}a and others},
	year         = 2021,
	journal      = {{arXiv}},
	doi          = {10.48550/arXiv.2105.07028}
}
@article{voss_full-stack_2017,
	title        = {{Full-stack genomics pipelining with GATK4 + WDL + Cromwell}},
	author       = {Voss, Kate and Gentry, Jeff and Auwera, Geraldine Van Der},
	year         = 2017,
	journal      = {{F1000Research}},
	doi          = {10.7490/f1000research.1114631.1}
}
@article{di_tommaso_nextflow_2017,
	title        = {{Nextflow enables reproducible computational workflows}},
	author       = {Di Tommaso, Paolo and Chatzou, Maria and Floden, Evan W. and Barja, Pablo Prieto and Palumbo, Emilio and Notredame, Cedric},
	year         = 2017,
	journal      = {{Nature Biotechnology}},
	volume       = 35,
	number       = 4,
	pages        = {316--319},
	doi          = {10.1038/nbt.3820}
}
@article{koster_snakemake_2012,
	title        = {{Snakemake---a scalable bioinformatics workflow engine}},
	author       = {K\"{o}ster, Johannes and Rahmann, Sven},
	year         = 2012,
	journal      = {{Bioinformatics}},
	volume       = 28,
	number       = 19,
	pages        = {2520--2522},
	doi          = {10.1093/bioinformatics/bts480}
}
@article{leprevost_biocontainers_2017,
	title        = {{BioContainers: An open-source and community-driven framework for software standardization}},
	author       = {Leprevost, Felipe da Veiga and Gr\"{u}ning, Bj\"{o}rn A. and Alves Aflitos, Saulo and R\"{o}st, Hannes L. and Uszkoreit, Julian and Barsnes, Harald and others},
	year         = 2017,
	journal      = {{Bioinformatics}},
	volume       = 33,
	number       = 16,
	pages        = {2580--2582},
	doi          = {10.1093/bioinformatics/btx192}
}
@article{soiland-reyes_packaging_2022,
	title        = {{Packaging research artefacts with RO-Crate}},
	author       = {Soiland-Reyes, Stian and Sefton, Peter and Crosas, Merc\`{e} and Castro, Leyla Jael and Coppens, Frederik and Fern\'{a}ndez, Jos\'{e} M. and others},
	year         = 2022,
	journal      = {{Data Science}},
	volume       = 5,
	number       = 2,
	pages        = {97--138},
	doi          = {10.3233/ds-210053}
}
@article{khan_sharing_2019,
	title        = {{Sharing interoperable workflow provenance: A review of best practices and their practical application in CWLProv}},
	author       = {Khan, Farah Zaib and Soiland-Reyes, Stian and Sinnott, Richard O. and Lonie, Andrew and Goble, Carole and Crusoe, Michael R.},
	year         = 2019,
	journal      = {{GigaScience}},
	volume       = 8,
	number       = 11,
	pages        = {giz095},
	doi          = {10.1093/gigascience/giz095}
}
@article{goble_implementing_2021,
	title        = {{Implementing FAIR digital objects in the EOSC-Life workflow collaboratory}},
	author       = {Goble, Carole and Soiland-Reyes, Stian and Bacall, Finn and Owen, Stuart and Williams, Alan and Eguinoa, Ignacio and others},
	year         = 2021,
	journal      = {{Zenodo}},
	publisher    = {Zenodo},
	doi          = {10.5281/zenodo.4605654}
}
@article{oconnor_dockstore_2017,
	title        = {{The Dockstore: Enabling modular, community-focused sharing of Docker-based genomics tools and workflows}},
	author       = {O'Connor, Brian D. and Yuen, Denis and Chung, Vincent and Duncan, Andrew G. and Liu, Xiang Kun and Patricia, Janice and others},
	year         = 2017,
	journal      = {{F1000Research}},
	volume       = 6,
	pages        = 52,
	doi          = {10.12688/f1000research.10137.1}
}
@article{ewels_nf-core_2020,
	title        = {{The nf-core framework for community-curated bioinformatics pipelines}},
	author       = {Ewels, Philip A. and Peltzer, Alexander and Fillinger, Sven and Patel, Harshil and Alneberg, Johannes and Wilm, Andreas and others},
	year         = 2020,
	journal      = {{Nature Biotechnology}},
	volume       = 38,
	number       = 3,
	pages        = {276--278},
	doi          = {10.1038/s41587-020-0439-x}
}
@article{baker_1500_2016,
	title        = {{1,500 scientists lift the lid on reproducibility}},
	author       = {Baker, Monya},
	year         = 2016,
	journal      = {{Nature}},
	volume       = 533,
	number       = 7604,
	pages        = {452--454},
	doi          = {10.1038/533452a}
}
@article{suetake_sapporo_2022,
	title        = {{Sapporo: A workflow execution service that encourages the reuse of workflows in various languages in bioinformatics}},
	author       = {Suetake, Hirotaka and Tanjo, Tomoya and Ishii, Manabu and P. Kinoshita, Bruno and Fujino, Takeshi and Hachiya, Tsuyoshi and others},
	year         = 2022,
	journal      = {{F1000Research}},
	volume       = 11,
	pages        = 889,
	doi          = {10.12688/f1000research.122924.1}
}
@misc{patel_rnaseq_2022,
	title        = {{nf-core/rnaseq: nf-core/rnaseq v3.7}},
	author       = {Harshil Patel and Phil Ewels and Alexander Peltzer and Rickard Hammar\'{e}n and Olga Botvinnik and Gregor Sturm and others},
	year         = 2022,
	publisher    = {Zenodo},
	doi          = {10.5281/zenodo.6513815}
}
@article{danecek_twelve_2021,
	title        = {{Twelve years of SAMtools and BCFtools}},
	author       = {Danecek, Petr and Bonfield, James K. and Liddle, Jennifer and Marshall, John and Ohan, Valeriu and Pollard, Martin O. and others},
	year         = 2021,
	journal      = {{GigaScience}},
	volume       = 10,
	number       = 2,
	pages        = {giab008},
	doi          = {10.1093/gigascience/giab008}
}
@article{rehm2021ga4gh,
	title        = {{GA4GH: International policies and standards for data sharing across genomic research and healthcare}},
	author       = {Rehm, Heidi L. and Page, Angela J.H. and Smith, Lindsay and Adams, Jeremy B. and Alterovitz, Gil and Babb, Lawrence J. and others},
	year         = 2021,
	journal      = {{Cell Genomics}},
	publisher    = {Elsevier},
	volume       = 1,
	number       = 2,
	pages        = 100029
}
@article{ison_edam_2013,
	title        = {{EDAM: An ontology of bioinformatics operations, types of data and identifiers, topics and formats}},
	author       = {Ison, Jon and Kalas, Mat\'{u}\v{s} and Jonassen, Inge and Bolser, Dan and Uludag, Mahmut and {McWilliam}, Hamish and others},
	year         = 2013,
	journal      = {{Bioinformatics}},
	volume       = 29,
	number       = 10,
	pages        = {1325--1332},
	doi          = {10.1093/bioinformatics/btt113}
}
@article{danecek_variant_2011,
	title        = {{The variant call format and VCFtools}},
	author       = {Danecek, Petr and Auton, Adam and Abecasis, Goncolo and Albers, Cornelis A. and Banks, Eric and DePristo, Mark A. and others},
	year         = 2011,
	journal      = {{Bioinformatics}},
	volume       = 27,
	number       = 15,
	pages        = {2156--2158},
	doi          = {10.1093/bioinformatics/btr330}
}
@article{suetake_workflow_2022,
	title        = {{Workflow sharing with automated metadata validation and test execution to improve the reusability of published workflows}},
	author       = {Suetake, Hirotaka and Fukusato, Tsukasa and Igarashi, Takeo and Ohta, Tazro},
	year         = 2022,
	journal      = {{bioRxiv}},
	doi          = {10.1101/2022.07.08.499265}
}
@misc{broad_gatk_2021,
	title        = {{broadinstitute/gatk - scripts/mitochondria\_m2\_wdl}},
	author       = {Peterson, Anders and Verdier, Axel and Abdel Ghany, Ayman and Bimber, Ben and Yoo, Byunggil and G\'{o}mez-S\'{a}nchez, Daniel and others},
	year         = 2021,
	publisher    = {GitHub},
	url          = {https://github.com/broadinstitute/gatk/tree/33bda5e08b6a09b40a729ee525d2e3083e0ecdf8/scripts/mitochondria\_m2\_wdl},
	howpublished = {\url{https://github.com/broadinstitute/gatk/tree/33bda5e08b6a09b40a729ee525d2e3083e0ecdf8/scripts/mitochondria\_m2\_wdl}}
}
@article{kodama_ddbj_2015,
	title        = {{The DDBJ Japanese Genotype-phenotype Archive for genetic and phenotypic human data}},
	author       = {Kodama, Yuichi and Mashima, Jun and Kosuge, Takehide and Katayama, Toshiaki and Fujisawa, Takatomo and Kaminuma, Eli and others},
	year         = 2015,
	journal      = {{Nucleic Acids Research}},
	publisher    = {Oxford University Press},
	volume       = 43,
	number       = {D1},
	pages        = {D18--D22}
}
@misc{suetake_tonkaz-test-methods_2022,
	title        = {{sapporo-wes/tonkaz - tests/README.md}},
	author       = {Suetake, Hirotaka},
	year         = 2022,
	publisher    = {GitHub},
	doi          = {10.5281/zenodo.7102376},
	url          = {https://github.com/sapporo-wes/tonkaz/blob/main/tests/README.md},
	howpublished = {\url{https://github.com/sapporo-wes/tonkaz/blob/main/tests/README.md}},
	version      = {0.2.3}
}
@misc{suetake_tonkaz-raw-data_2022,
	title        = {{Raw data of workflow execution results used in Tonkaz's experiments}},
	author       = {Suetake, Hirotaka},
	year         = 2022,
	publisher    = {Zenodo},
	doi          = {10.5281/zenodo.7098337},
	version      = {1.0.0}
}
@article{mckenna_genome_2010,
	title        = {{The Genome Analysis Toolkit: A MapReduce framework for analyzing next-generation DNA sequencing data}},
	author       = {{McKenna}, Aaron and Hanna, Matthew and Banks, Eric and Sivachenko, Andrey and Cibulskis, Kristian and Kernytsky, Andrew and others},
	year         = 2010,
	journal      = {{Genome Research}},
	volume       = 20,
	number       = 9,
	pages        = {1297--1303},
	doi          = {10.1101/gr.107524.110},
	url          = {http://genome.cshlp.org/lookup/doi/10.1101/gr.107524.110}
}
@article{dobin_star_2013,
	title        = {{STAR: Ultrafast universal RNA-seq aligner}},
	author       = {Dobin, Alexander and Davis, Carrie A. and Schlesinger, Felix and Drenkow, Jorg and Zaleski, Chris and Jha, Sonali and others},
	year         = 2013,
	journal      = {{Bioinformatics}},
	volume       = 29,
	number       = 1,
	pages        = {15--21},
	doi          = {10.1093/bioinformatics/bts635}
}
@article{pertea_stringtie_2015,
	title        = {{StringTie enables improved reconstruction of a transcriptome from RNA-seq reads}},
	author       = {Pertea, Mihaela and Pertea, Geo M. and Antonescu, Corina M. and Chang, Tsung-Cheng and Mendell, Joshua T. and Salzberg, Steven L.},
	year         = 2015,
	journal      = {{Nature Biotechnology}},
	volume       = 33,
	number       = 3,
	pages        = {290--295},
	doi          = {10.1038/nbt.3122}
}
@misc{nf-core_test-datasets_2018,
	title        = {{nf-core/test-datasets - rnaseq/samplesheet/v3.4/samplesheet\_test.csv}},
	author       = {{nf-core Community}},
	year         = 2018,
	publisher    = {GitHub},
	url          = {https://raw.githubusercontent.com/nf-core/test-datasets/rnaseq/samplesheet/v3.4/samplesheet\_test.csv},
	howpublished = {\url{https://raw.githubusercontent.com/nf-core/test-datasets/rnaseq/samplesheet/v3.4/samplesheet\_test.csv}}
}
@misc{suetake_small-dataset_2022,
	title        = {{sapporo-wes/test-workflow: 1.0.2 - assets/nf-core\_rnaseq\_samplesheet\_small\_test.csv}},
	author       = {Suetake, Hirotaka and Ohta, Tazro},
	year         = 2022,
	publisher    = {Zenodo},
	doi          = {10.5281/zenodo.7102664},
	url          = {https://raw.githubusercontent.com/sapporo-wes/test-workflow/1.0.2/assets/nf-core\_rnaseq\_samplesheet\_small\_test.csv},
	version      = {1.0.2},
	howpublished = {\url{https://raw.githubusercontent.com/sapporo-wes/test-workflow/1.0.2/assets/nf-core\_rnaseq\_samplesheet\_small\_test.csv}}
}
@article{ewels_multiqc_2016,
	title        = {{MultiQC: Summarize analysis results for multiple tools and samples in a single report}},
	author       = {Ewels, Philip and Magnusson, M\aa{}ns and Lundin, Sverker and K\"{a}ller, Max},
	year         = 2016,
	journal      = {{Bioinformatics}},
	volume       = 32,
	number       = 19,
	pages        = {3047--3048},
	doi          = {10.1093/bioinformatics/btw354}
}
@misc{suetake_test-workflow_2022,
	title        = {{sapporo-wes/test-workflow: 1.0.2}},
	author       = {Suetake, Hirotaka and Ohta, Tazro},
	year         = 2022,
	publisher    = {Zenodo},
	doi          = {10.5281/zenodo.7102664},
	version      = {1.0.2}
}
@article{nust_codecheck_2021,
	title = {{{CODECHECK}: An Open Science initiative for the independent execution of computations underlying research articles during peer review to improve reproducibility}},
	volume = {10},
	doi = {10.12688/f1000research.51738.1},
	pages = {253},
	journal = {F1000Research},
	author = {Nüst, Daniel and Eglen, Stephen J.},
	year = {2021},
}
