## Supplementary material for "A workflow reproducibility scale for automatic validation of biological interpretation results": LaTeX Source: main.tex

\documentclass[times, twoside, watermark]{zHenriquesLab-StyleBioRxiv}

\usepackage{blindtext}
\usepackage{graphicx}
\usepackage{siunitx}
\usepackage{flushend}
\usepackage{url}
\usepackage{booktabs}
\usepackage{multirow}
\usepackage{tabularx}

\leadauthor{Suetake}

\begin{document}

\title{A workflow reproducibility scale for automatic validation of biological interpretation results}
\shorttitle{Workflow reproducibility scale}

\author[1]{Hirotaka Suetake}  %
\author[2]{Tsukasa Fukusato}  %
\author[1]{Takeo Igarashi}  %
\author[3,\Letter]{Tazro Ohta}

\affil[1]{Department of Creative Informatics, Graduate School of Information Science and Technology, The University of Tokyo, Tokyo, Japan}
\affil[2]{Department of Computer Science, Graduate School of Information Science and Technology, The University of Tokyo, Tokyo, Japan}
\affil[3]{Database Center for Life Science, Joint Support-Center for Data Science Research, Research Organization of Information and Systems, Shizuoka, Japan}

\maketitle

\begin{abstract}
    \textbf{\textit{Background:}}
    Reproducibility of data analysis workflow is a key issue in the field of bioinformatics.
    Recent computing technologies, such as virtualization, have made it possible to reproduce workflow execution with ease.
    However, the reproducibility of results is not well discussed; that is, there is no standard way to verify whether the biological interpretation of reproduced results are the same.
    Therefore, it still remains a challenge to automatically evaluate the reproducibility of results.

    \vspace{1.5mm}
    \textbf{\textit{Results:}}
    We propose a new metric, a reproducibility scale of workflow execution results, to evaluate the reproducibility of results.
    This metric is based on the idea of evaluating the reproducibility of results using biological feature values (e.g., number of reads, mapping rate, and variant frequency) representing their biological interpretation.
    We also implemented a prototype system that automatically evaluates the reproducibility of results using the proposed metric.
    To demonstrate our approach, we conducted an experiment using workflows used by researchers in real research projects and the use cases that are frequently encountered in the field of bioinformatics.

    \vspace{1.5mm}
    \textbf{\textit{Conclusions:}}
    Our approach enables automatic evaluation of the reproducibility of results using a fine-grained scale.
    By introducing our approach, it is possible to evolve from a binary view of whether the results are superficially identical or not to a more graduated view.
    We believe that our approach will contribute to more informed discussion on reproducibility in bioinformatics.

\end{abstract}

\begin{keywords}
    Workflow | Provenance | Reproducibility
\end{keywords}

\begin{corrauthor}
   
\end{corrauthor}

\section*{Background}

Bioinformatics is big data science and is considered the most demanding domain in terms of data acquisition, storage, distribution, and analysis~\citep{stephens_big_2015}.
Because the low cost and high throughput of measurement instruments have made it possible to generate large amounts of data, large-scale data analysis using a computer is required to extract valuable knowledge from the data~\citep{stein_case_2010, goodwin_coming_2016}.
For each data analysis process, such as data transformation, public database referencing and merging, and statistical processing, much open-source software is developed and released~\citep{prins_toward_2015}.
Researchers typically choose appropriate software for each analysis process, build a workflow by combining the software, and execute the workflow in a computing environment~\citep{perkel_workflow_2019}.
However, it can be challenging to ensure the reproducibility of data analysis due to a number of factors, such as a large amount of data, the diversity of data types and software, and the complexity of the computing environment~\citep{banati_evaluating_2016}.

Reproducibility of research is an essential issue in the scientific community~\citep{noauthor_software_2014, noauthor_rebooting_2015}.
However, Baker raised the alarm of a ``reproducibility crisis'' based on survey results that \textit{``more than 70\% of researchers have tried and failed to reproduce another scientist's experiments, and more than half have failed to reproduce their own experiments''}~\citep[p.~452]{baker_1500_2016}.
The key here is the requirement for research to be considered reproducible.
Drummond argued that replicability and reproducibility are often confused, but they are different concepts and need to be clearly distinguished~\citep{drummond_replicability_2009}.
The Association for Computing Machinery (ACM) also attempts to define the terms repeatability, reproducibility, and replicability (Table~\ref{table:acmDefinitions})~\citep{acm_artifact_2021}.
While these definitions are in the context of computerized analysis, it should be noted that most existing studies have focused on whether the execution can be reproduced or not and have not considered the verification of the results.
That is, they only state that the resulting data are exactly the same as in the original but do not adequately discuss the verification of whether the results are reproducible or not.
Therefore, the reproducibility of data analysis can be divided into two parts: the execution of the analysis and the verification of the results. We will focus our discussion on the second part, verification.

\begin{table*}[ht!]
    \centering
    \caption{
        Repeatability, reproducibility, and replicability.
        According to the ACM~\citep{acm_artifact_2021}, repeatability is defined as a researcher can reliably repeat her own computation.
        Reproducibility is defined as an independent group can obtain the same result using the author's own artifacts, and replicability is defined as an independent group can obtain the same result using artifacts which they develop completely independently.
    }
    \label{table:acmDefinitions}
    \begin{tabular}{lllll}
        \toprule
        Term            & Team      & Environment & Setup (code and data) & Result \\
        \midrule
        Repeatability   & Same      & Same        & Same                  & Same   \\
        Reproducibility & Different & Different   & Same                  & Same   \\
        Replicability   & Different & Different   & Different             & Same   \\
        \bottomrule
    \end{tabular}
\end{table*}

Many workflow systems have been developed to improve the efficiency of building and executing complex data analysis~\citep{leprevost_best_2014, wratten_reproducible_2021, amstutz_existing_2021}.
Each system has unique characteristics, but in particular, workflow systems can have a syntax for describing the data analysis, called a workflow language.
Large user communities have been formed around these workflow languages.
The Common Workflow Language (CWL)~\citep{crusoe_methods_2021}, the Workflow Description Language (WDL)~\citep{voss_full-stack_2017}, Nextflow~\citep{di_tommaso_nextflow_2017}, and Snakemake~\citep{koster_snakemake_2012} are typical examples.
These systems also have execution systems that work with computational frameworks, such as job schedulers, container runtimes, and package managers.
Thus, these workflow systems facilitate the execution of data analysis by different teams and in different environments through the use of virtualization technology and syntax that abstracts software and computational requirements~\citep{leprevost_biocontainers_2017}.

The advent and widespread use of workflow systems have facilitated data analysis re-execution.
However, as mentioned earlier, to ensure reproducibility, it is necessary to verify the execution results, that is, whether the same biological interpretation is obtained or not.
To address this issue, frameworks, such as Research Object Crate (RO-Crate)~\citep{soiland-reyes_packaging_2022} and CWLProv~\citep{khan_sharing_2019}, have been proposed to generate workflow provenance, a structured archive that packages workflow-related metadata, such as workflow descriptions, execution parameters, input and output data, tests, and documentation, in a machine-readable format.
This provenance information is distributed on workflow sharing platforms, such as WorkflowHub~\citep{goble_implementing_2021}, Dockstore~\citep{oconnor_dockstore_2017}, and nf-core~\citep{ewels_nf-core_2020}.
When appropriate provenance is provided by the author, a researcher can use this information to verify new execution results making the process reproducible.

However, the process of comparing the provenance and execution results is often incomplete and inefficient.
In automatic comparison, the checksums of the output files are used; however, they do not always match.
This is because these checksums may differ depending on the software version, timestamps, heuristic algorithms, and computing environments (e.g., OS and CPU architecture, etc.).
However, the same biological interpretation may be obtained even when the output files do not match exactly; for example, only the timestamps in the output files may differ.
Thus, a simple comparison using a checksum is incomplete in verifying results.
Another method is to have humans semantically interpret the results.
However, due to its inefficiency, this method is not possible when the scale of the data analysis execution is large.
From the above, the verification of results using provenance remains challenging because the current procedure is limited to incomplete automatic comparison and inefficient manual checking.

Automation is essential for the verification of practical workflows that output many files; however, binary determination by checking checksums is not sufficient.
Thus, it is necessary to introduce a fine-grained scale to determine the degree of reproducibility of the results.
Automatic verification of results using this scale will make verification of workflow reproducibility practical.
In this paper, we propose a reproducibility scale of workflow execution results based on some discussion and experiments and a validation method using this scale.
We implement a workflow execution system that generates a workflow provenance that contains metadata required for verification.
This implementation is an extension of Sapporo~\citep{suetake_sapporo_2022}, an existing workflow execution service (WES).
Sapporo's extensibility makes it compatible with various workflow languages and execution systems.
In addition, we adapt RO-Crate as the workflow provenance format.
We also develop Tonkaz: a command-line tool that verifies the reproducibility of data analysis results by comparing the workflow provenances.
To demonstrate the effectiveness of our approach, we apply it to workflows used by researchers in real research projects.
The full reproducibility of research is still an issue that has not been fully resolved.
Nevertheless, we hope our approach will contribute to solving this problem by increasing the resolution of the definition of reproducibility.

\section*{Methods}

\subsection*{Reproducibility scale of workflow execution results}

A workflow is a sequence of computational steps that combine analysis tools according to their inputs and outputs.
The first tool takes input data and passes its output on as input for the next tool.
Thus, the result of the workflow execution is the cumulative output of each tool or the last tool in the workflow.
It should be noted that the output of a tool includes not only output files, but also execution logs (e.g., standard output and error) and runtime information (e.g., exit code, start time, and end time).
Returning to the purpose of data analysis here, it is to obtain useful biological knowledge from the data.
Therefore, it is not sufficient to consider the output files and logs as the only result of the workflow execution; the biological feature values interpreted from the output files and logs should be considered as the result of the workflow execution.

The format to represent biological features obtained from data analysis is not standardized and varies depending on the analysis tool.
For example, there are summarized formats (tabular and graph) and formats that express biological features themselves, such as Sequence Alignment/Map (SAM) and the Variant Call Format (VCF).
To interpret and verify the results, the individual executing the workflow visually checks the output graph or uses a tool to extract a numerical feature value from the file, for example, SAMtools~\citep{danecek_twelve_2021} to extract mapping statistics from the SAM format.
Because these processes require domain knowledge, it is ideal that the workflow itself provides a structured summary and a way to interpret it.
However, this depends on the skill and effort of the individual workflow developer, and the diversity of tools and workflows makes it challenging to provide them in a standardized way.

There are several workflows that provide a way to verify reproducibility using biological feature values.
For example, the RNA-seq workflow~\citep{patel_rnaseq_2022} distributed by the nf-core project has a test mode to verify that the workflow is working as expected.
In this mode, the workflow is executed with a small test dataset, and the biological feature values are compared with the expected values.
The mapping rate, which represents the percentage of reads that are mapped to the reference genome, is used as a biological feature value.
If the difference between the values is within the threshold, the workflow is considered to be working as expected.
As a preliminary experiment, we compared the output files without using such biological feature values---that is, the checksum method was used to verify an exact match of a file.
As a result, when we executed the above RNA-seq workflow twice in the same environment and compared the file output BAM files (the compressed binary version of the SAM files), we found that the checksum values were different and the file sizes differed by several bytes (see the Section ``Results'' for details).
It is ideal that the output files are exactly the same, but it is difficult to achieve this goal because the output files are generated by the analysis tools, and these tools are not designed to produce the same output all the time.
Therefore, we concluded that it is not sufficient to check only the exact match of output files to verify the reproducibility of workflow execution results and that a method using biological feature values and threshold should also be introduced.

Based on the above discussion and preliminary experiments, we propose a method to verify the reproducibility of workflow execution results using biological feature values and threshold.
The method consists of two steps: (1) extracting biological feature values from the output files and logs and (2) comparing the extracted biological feature values with the expected values using threshold values.
A detailed description of each step is provided in the Sections ``Generation of workflow provenance containing biological feature values'' and ``Automatic verification of reproducibility.''
We also propose a scale to evaluate the reproducibility of workflow execution results based on the method (Table~\ref{tab:reproducibilityScale}).
This allows the reproducibility of results to be expressed at a higher resolution than a binary measure of whether the results are the same or not.

\begin{table*}[ht!]
    \centering
    \caption{
        Reproducibility scale of workflow execution results.
        For each of the output files, this is determined by comparing the expected provenance with the provenance of the actual execution.
        If the file of the same name in the expected provenance and the actual execution are identical, the file is considered to be fully reproduced.
        If the file of the same name is not identical, it is determined whether its difference is acceptable or not using the feature and threshold values.
        If the difference is acceptable, the file is considered to be partially reproduced.
        If the file exists in the expected provenance but not in the actual execution, it is considered to be not reproduced.
    }
    \label{tab:reproducibilityScale}
    \begin{tabularx}{\textwidth}{lcX}
        \toprule
        Reproducibility scale    & Level & Description                                                                                                                                                        \\
        \midrule
        Fully Reproduced         & 3     & Output files are identical with the same checksum.                                                                                                                 \\
        Acceptable Differences   & 2     & Output files are not identical, but their biological feature values (e.g., number of reads, mapping rate, and variant frequency) are similar (within a threshold). \\
        Unacceptable Differences & 1     & Output files are not identical, and their biological feature values are not similar (beyond threshold).                                                            \\
        Not Reproduced           & 0     & The workflow does not produce output files.                                                                                                                        \\
        \bottomrule
    \end{tabularx}
\end{table*}

\subsection*{Generation of workflow provenance containing biological feature values}

To verify the reproducibility of workflow execution results using biological feature values, it is necessary to package the workflow execution results as the workflow provenance in a standardized format.
Because there are many workflow languages and execution engines, we first abstracted the workflow execution itself.
Thus, we extended Sapporo, an existing WES implementation.
Sapporo has an API scheme that satisfies the Global Alliance for Genomics and Health (GA4GH) Workflow Execution Service (WES) standard~\cite{rehm2021ga4gh}, enabling the workflow execution and results acquisition in a standardized manner.
In addition, due to its extensibility, it can execute workflows written in various languages, such as CWL, WDL, Nextflow, and Snakemake.
Therefore, by extending Sapporo, workflow execution written in various languages can be handled in the same way.

When a workflow is executed in Sapporo, the files related to the execution are stored in the file system as workflow provenance.
This provenance directory contains the workflow definition files, input files, intermediate files, output files, log files, execution parameters, runtime information, etc.
Thus, we converted Sapporo's provenance into RO-Crate, a standardized format for packaging research objects expressed in JSON-LD.
Because the RO-Crate use case included the packaging of workflow execution results, it was sufficient to map Sapporo's provenance to the ontology provided by RO-Crate.
However, for verification, we defined some additional terms and properties\footnote{\url{https://raw.githubusercontent.com/sapporo-wes/sapporo-service/main/sapporo/ro-terms.csv}}.
For example, we defined the property ``mappedRate'' to represent the mapping rate of the output file, which is a biological feature value used for verification.
In addition, RO-Crate is designed to rely on the local file system for file location resolution and checksum representation.
However, we prioritized the portability of being able to carry the provenance in a single file, so we put all the information necessary for verification, such as checksums, biological feature values, and contents of files of small size, in the RO-Crate file.

Because the workflow output freely produces a large number of diverse files, it is impractical to extract biological feature values for all files.
Thus, we used the file extension to determine the file type and used an appropriate tool to extract the biological feature values.
We used the file types defined in the EDAM ontology~\citep{ison_edam_2013}, which are widely used to express biological interpretations (Table~\ref{tab:edamFileTypes}).
For example, if the file type is SAM, we used SAMtools to extract the number of reads and the mapping rate, and if the file type is VCF, we used VCFtools~\citep{danecek_variant_2011} to extract the number of variants and the variant frequency.
In addition, the number of lines in the file is also a biological feature value.
For example, if the file type is FASTQ, four lines represent one sequence read.
Figure~\ref{fig:exampleFileEntity} is an example of a file entity in RO-Crate, which contains the biological feature values, such as the statistics obtained from the file, file size, and the number of lines.

\begin{table*}[ht!]
    \centering
    \vspace{\baselineskip}
    \caption{
        File types and extensions defined in EDAM ontology.
        These file types and extensions are used to extract biological feature values from the output files.
    }
    \label{tab:edamFileTypes}
    \begin{tabular}{ccl}
        \toprule
        EDAM ID               & File type & Extension                               \\
        \midrule
        \texttt{format\_1929} & FASTA     & \texttt{.fa, .fasta}                    \\
        \texttt{format\_1930} & FASTQ     & \texttt{.fq, .fastq, .fq.gz, .fastq.gz} \\
        \texttt{format\_1975} & GFF       & \texttt{.gff, .gff3}                    \\
        \texttt{format\_2306} & GTF       & \texttt{.gtf}                           \\
        \texttt{format\_2572} & BAM       & \texttt{.bam}                           \\
        \texttt{format\_2573} & SAM       & \texttt{.sam}                           \\
        \texttt{format\_3003} & Bed       & \texttt{.bed}                           \\
        \texttt{format\_3004} & BigBed    & \texttt{.bb}                            \\
        \texttt{format\_3005} & Wig       & \texttt{.wig}                           \\
        \texttt{format\_3006} & BigWig    & \texttt{.bw}                            \\
        \texttt{format\_3016} & VCF       & \texttt{.vcf, .vcf.gz}                  \\
        \bottomrule
    \end{tabular}
\end{table*}

\begin{figure*}[ht!]
    \centering
    \includegraphics[width=\textwidth]{figures/example_file_entity.pdf}
    \caption{
        An example of a file entity in RO-Crate.
        This is a part of the actual workflow execution results and uses the RNA-seq workflow distributed by the nf-core project.
        The output file in this example is a BAM file, and its biological feature values are the file size, number of mapped reads, and mapping rate.
        Thus, the file entity contains properties such as ``contentSize,'' ``stats:mappedReads,'' and ``stats:mappedRate''.
        These are defined as additional terms in Sapporo, and the values are extracted from the file using SAMtools.
    }
    \label{fig:exampleFileEntity}
\end{figure*}

For provenance enrichment and sharing, we integrated Sapporo and Yevis~\citep{suetake_workflow_2022}.
Yevis is a system that builds a workflow registry and also acts as a client for Sapporo.
The workflow metadata file used in Yevis contains not only the information required to execute the workflow but also the information for workflow availability and traceability in workflow sharing, such as author, open-source license, and documentation link.
Thus, by executing the workflow in Sapporo via Yevis, the availability and traceability of the generated provenance are improved.
In addition, because Yevis' workflow sharing feature enables the attachment of generated provenance to shared workflows, the reproducibility of shared workflows can be verified by other users.

From the above, by executing the workflow with Sapporo and Yevis (Figure~\ref{fig:tonkazOverview}), a workflow provenance containing feature values representing a biological interpretation is generated as RO-Crate.
This method also applies to workflows written in various languages and can address a wide range of use cases, such as workflow sharing.
Therefore, by generating and sharing a provenance containing biological feature values, it is possible to verify the reproducibility of the workflow execution results in other users' environments.

\subsection*{Automatic verification of reproducibility}

We developed Tonkaz to automatically verify the reproducibility of workflow execution results by comparing the biological feature values contained in the workflow provenance.
One use case of Tonkaz is to compare the expected result, which is provided by a workflow developer, and the actual result, which is generated in the user's environment (Figure~\ref{fig:tonkazOverview}).
That is, Tonkaz verifies that the results are the same according to the ACM's definition of reproducibility (Table~\ref{table:acmDefinitions}).
Another use case is ACM's definition of repeatability, which is to verify that the results are the same even if a workflow is executed multiple times in the same environment, and it will not be broken by updates to dependencies.
Thus, these use cases indicate that we must verify the reproducibility of the results, regardless of the differences in execution methods and environments.

\begin{figure*}[ht!]
    \centering
    \includegraphics[width=0.8\textwidth]{figures/tonkaz_overview.pdf}
    \caption{
        The flowchart representing the Tonkaz use case.
        The workflow built by the workflow developer is executed by WES, which is a combination of Sapporo and Yevis, and the workflow provenance, including feature values of the output files, is generated in RO-Crate format.
        This provenance is used as the expected value for the verification of reproducibility.
        Using the shared workflow, the user executes the shared workflow in his/her own environment using WES.
        Using Tonkaz, the user then compares the shared provenance with the provenance generated by the user's workflow execution and verifies the reproducibility of the workflow execution results.
    }
    \vspace{\baselineskip}
    \vspace{\baselineskip}
    \label{fig:tonkazOverview}
\end{figure*}

We designed Tonkaz to accept as arguments two RO-Crates, one containing the expected provenance and the other containing the actual provenance.
Tonkaz compares the biological feature values of the output files in the two RO-Crates and calculates the reproducibility scale for each file.
The files to be compared are those output files that have been assigned EDAM ontology, as described in Table~\ref{tab:edamFileTypes}.
This is because the workflow output files often include log files that are not related to the biological interpretation and image files that are not mechanically comparable.
For example, nf-core's RNA-seq workflow produces 872 files, but only 25 files are assigned EDAM ontology.
In the process of comparing files and calculating the reproducibility scale, Tonkaz first checks whether the files are identical using a checksum (Figure~\ref{fig:tonkazReproScale}).
If the files are identical, the reproducibility scale is ``Fully Reproduced.''
If the files are not identical, Tonkaz compares the biological feature values of the files using a threshold value to determine whether the differences are acceptable or not.
The default threshold value used is 0.05, but this value can be changed according to the use case.
This is because some workflows for medical applications or quality control of biological materials require a lower threshold value.
The comparison result is finally summarized in a table, and the reproducibility scale of each file is also displayed in a table (Figure~\ref{fig:tonkazCliExample}).
However, Tonkaz does not score the reproducibility of the entire workflow.
This is because, again, the purpose of comparison may differ depending on the use case, and it is not practical to automate the final decision.
Thus, we implemented an option to generate structured data in Tonkaz.
In addition, we believe that workflow developers should use this option and write conditions or scripts to determine the reproducibility for each use case.

\begin{figure*}[ht!]
    \centering
    \includegraphics[width=0.7\textwidth]{figures/tonkaz_repro_scale.pdf}
    \caption{
        The process for calculating the reproducibility scale of a file.
        Tonkaz first checks whether the files are identical using a checksum.
        If the files are identical, the reproducibility scale value is ``Fully Reproduced.''
        If the files are not identical, Tonkaz compares the biological feature values of the files using a threshold value to determine whether the differences are acceptable or not.
        If the differences are acceptable, the reproducibility scale value is ``Acceptable Difference.''
        If the differences are unacceptable, the reproducibility scale value is ``Unacceptable Difference.''
        The default threshold value used is 0.05, but this value can be changed according to the use case.
        If the file entity exists only in one of the two RO-Crates, the reproducibility scale value is ``Not Reproduced.''
    }
    \label{fig:tonkazReproScale}
\end{figure*}

\begin{figure*}[ht!]
    \centering
    \includegraphics[width=0.85\textwidth]{figures/tonkaz_cli_example.pdf}
    \caption{
        Example of the Tonkaz output.
        Tonkaz displays a table for each file and a final summary table.
        The user checks those summary tables to determine the reproducibility of the entire workflow and the differences between the expected and actual files (e.g., by using the diff command).
    }
    \label{fig:tonkazCliExample}
\end{figure*}

\section*{Results}

To demonstrate the effectiveness of our approach, we verified the reproducibility of workflow execution results by comparing the results of public workflows used by researchers in real research projects, not simple ones for testing.
This verification was based on the following five practical use cases: (1) execution in the same environment, (2) execution in a different environment, (3) execution of different versions of the workflow, (4) execution with missing datasets, and (5) comparison using all output files.
We used the following three workflows: (1) the mitochondrial short variant discovery workflow distributed a GATK best practice workflow (hereafter referred to as GATK workflow, language: WDL)~\citep{broad_gatk_2021}, (2) RNA-Seq workflow distributed by nf-core (hereafter referred to as RNA-seq workflow, language: Nextflow)~\citep{patel_rnaseq_2022}, and (3) GATK best practice-compatible germline short variant discovery workflow, which is used to process whole-genome sequencing data of the Japanese Genotype-phenotype Archive (hereafter referred to as JGA workflow, language: CWL)~\citep{kodama_ddbj_2015}.
We used the following two execution environments: (1) Ubuntu 20.04 LTS (CPU: Intel Xeon E5-2640 @ 2.50GHz, RAM: 24GB, Docker: 20.10.8) and (2) macOS 12.5.1 (CPU: Apple M1 Max, RAM: 64GB, Docker: 20.10.16).
Table~\ref{tab:workflowExecutionTable} shows the setting for each execution as a combination, and Table~\ref{tab:comparisonResults} summarizes the verification results based on the use cases.
The methods and results of the workflow execution and verification are described in the online documentation ``sapporo-wes/tonkaz - tests/README.md''~\citep{suetake_tonkaz-test-methods_2022} and are published on Zenodo~\citep{suetake_tonkaz-raw-data_2022}.

\begin{table*}[ht!]
    \centering
    \caption{
        Combination table of workflow execution and execution settings.
        The first column is the definition of the execution name.
        In the second column and below are the workflow execution settings.
        The blank cells in the third and fourth columns indicate that there are no differences in the execution settings.
    }
    \label{tab:workflowExecutionTable}
    \begin{tabularx}{0.56\textwidth}{lcccc}
        \toprule
        Execution name & Workflow                 & Version & Dataset  & Environment \\
        \midrule
        GATK\_1st      & \multirow{3}{*}{GATK}    &         &          & Linux       \\
        GATK\_2nd      &                          &         &          & Linux       \\
        GATK\_mac      &                          &         &          & Mac         \\
        \hline
        JGA\_1st       & \multirow{3}{*}{JGA}     &         &          & Linux       \\
        JGA\_2nd       &                          &         &          & Linux       \\
        JGA\_mac       &                          &         &          & Mac         \\
        \hline
        RNA-seq\_1st   & \multirow{5}{*}{RNA-seq} & v3.7    & Standard & Linux       \\
        RNA-seq\_2nd   &                          & v3.7    & Standard & Linux       \\
        RNA-seq\_mac   &                          & v3.7    & Standard & Mac         \\
        RNA-seq\_v3.6  &                          & v3.6    & Standard & Linux       \\
        RNA-seq\_small &                          & v3.7    & Small    & Linux       \\
        \bottomrule
    \end{tabularx}
\end{table*}

\begin{table*}[ht!]
    \centering
    \vspace{\baselineskip}
    \caption{
        Comparisons of execution and verification results.
        The definition of each execution is defined in Table~\ref{tab:workflowExecutionTable}.
        Five use cases are assigned according to the combination of executions.
        In the fifth column and below are the number of files for each reproducibility scale defined in Table~\ref{tab:reproducibilityScale}: Level 3 is ``Fully Reproduced,'' Level 2 is ``Acceptable Difference,'' Level 1 is ``Unacceptable Difference,'' and Level 0 is ``Not Reproduced.''
    }
    \label{tab:comparisonResults}
    \begin{tabular}{clllcccc}
        \toprule
        ID & Source execution & Target execution & Use case              & Level 3 & Level 2 & Level 1 & Level 0 \\
        \midrule
        C1 & GATK\_1st        & GATK\_2nd        & Same environment      & 0       & 5       & 0       & 0       \\
        C2 & GATK\_1st        & GATK\_mac        & Different environment & 0       & 0       & 0       & 5       \\
        C3 & JGA\_1st         & JGA\_2nd         & Same environment      & 0       & 4       & 0       & 0       \\
        C4 & JGA\_1st         & JGA\_mac         & Different environment & 0       & 0       & 0       & 4       \\
        C5 & RNA-seq\_1st     & RNA-seq\_2nd     & Same environment      & 20      & 5       & 0       & 0       \\
        C6 & RNA-seq\_1st     & RNA-seq\_mac     & Different environment & 0       & 0       & 0       & 25      \\
        C7 & RNA-seq\_1st     & RNA-seq\_v3.6    & Different version     & 13      & 12      & 0       & 0       \\
        C8 & RNA-seq\_1st     & RNA-seq\_small   & Missing dataset       & 8       & 5       & 7       & 5       \\
        C9 & RNA-seq\_1st     & RNA-seq\_2nd     & All output files      & 557     & 306     & 1       & 8       \\
        \bottomrule
    \end{tabular}
\end{table*}

The Comparisons C1, C3, and C5 present execution results in the same environment.
Comparison C1 was performed using the GATK workflow, and the output file types were BAM and VCF.
The reproducibility scale value was Level 2 (Acceptable Difference) for all files, with no differences in biological feature values expressing biological interpretation (e.g., mapping rate and variant frequency).
The difference between these files was due to the fact that both the BAM and VCF files included the file paths of the original input file and timestamps in the header lines.
Thus, when using the analysis tool GATK~\citep{mckenna_genome_2010}, it is challenging to fully reproduce the output files because of the behavior that the output files contain the file paths and timestamps.
Comparison C3 was performed using the JGA workflow, and the output file types were VCF.
This result also showed no differences in biological feature values, and the differences in file contents were due to the behavior of GATK.
Comparison C5 was performed using the RNA-seq workflow, and the output file types were BAM, GTF, and BED.
All GTF and BED files were Level 3 (Fully Reproduced), and all BAM files were Level 2 (Acceptable Difference).
The difference between BAM files was due to the different order of the mapped reads in the BAM file.
These BAM files were mapped by STAR~\citep{dobin_star_2013}, and sorted by SAMtools; however, differences occurred.
These results show cases in which the output files were not identical, although the biological feature values were equal, due to the behavior of the analysis tool.

The Comparisons C2, C4, and C6 present the execution results in different environments.
All of these comparison results were Level 0 (Not Reproduced) because all execution in the Mac environment either failed or never finished.
All workflows used in this experiment were Docker containerized and were designed to be very reproducible in the execution context; however, runtime errors occurred due to the Arm processor architecture of the Mac environment.
Thus, even a very well-considered workflow may not be reproducible in a different environment.
In such cases, it is essential to increase the debuggability of the cause of the irreproducibility of the execution results.
Therefore, the importance of this debuggability indicates that it is helpful to include information about the execution environment in the workflow provenance; our approach and RO-Crate address them.

Comparison C7 presents the execution results in different versions.
Workflow developers often check for workflow breakage when updating versions of analysis tools included in the workflow.
In the RNA-seq workflow used in this comparison, the dependent analysis tools STAR, SAMtools, and StringTie~\citep{pertea_stringtie_2015} were updated with the workflow update from v3.6 to v3.7.
As a result of the comparison (C7), the number of files with Level 2 increased compared to C5, a comparison involving the same version.
The file types that became Level 2 were GTF, BED, and BAM; the GTF and BED files were newly changed from Level 3 to Level 2 when compared to C5.
The differences between the GTF files were due to differences in the FPKM field values and the version of StringTie included in the header line.
The BED files had a different number of lines, and the BAM files had a different number of mapped reads; however, those differences were within the threshold value.
This result indicates that verification using biological feature values and threshold is effective because apparent differences occur in output due to version updates and other reasons, and it is necessary to determine whether these differences are acceptable or not.

Comparison C8 presents the execution results in a case where the input dataset was partially missing.
The dataset used in RNA-seq\_1st contains six sequence read files (FASTQ)~\citep{nf-core_test-datasets_2018}, while the dataset used in RNA-seq\_small contained four sequence read files~\citep{suetake_small-dataset_2022}.
As a result of the comparison, the output files related to the sample with half the number of reads were Level 1 (Unacceptable Difference), while the sample with zero reads was Level 0.
In this case, setting the threshold used for verification to, for example, 0.5 instead of 0.05 (default value) will verify that the workflow is functioning as expected.
That is, this suggests that the threshold value and final decision may vary depending on the objectives of developers and users.

Comparison C9 presents the execution results in a case where all the output files were compared.
Most of the files were Level 3 or Level 2; however, 16 files were not reproduced (Level 0).
These level 0 files had random names or timestamps in the file names, for example, \texttt{mqc\_mqc\_mplplot\_gtnuqiebfc\_1.pdf} and \texttt{execution\_report\_2022-09-08\_06-28-19.html}.
Therefore, it is not appropriate to use all files to verify the reproducibility of execution results; it is essential to focus on characteristic files, such as BAM and GTF files.

For the five practical use cases, we found that our approach was well suited to verify the reproducibility of the workflow execution results.
In all use cases, existing methods that use checksums to verify exact file matches can produce false positives; this means that the workflow is considered not reproduced, even though it is working as expected.
Therefore, it is important to introduce a reproducibility scale and verify the workflow execution results' reproducibility at higher resolutions.

\section*{Discussion}

Despite its complexity, data analysis in bioinformatics is considered reproducible and is being shared.
In particular, the workflows shared by nf-core and GATK best practices are well maintained and include test datasets, documentation, and open-source licenses.
Ideally, all shared workflows would be like these; however, in reality, this is challenging because of the amount of work and domain knowledge required.
Thus, we aim to facilitate workflow sharing by providing a workflow provenance model and a workflow provenance verification method.
However, our approach is not applicable in domains where it is difficult to verify the results and inferences using a computer.
In such cases, it is first necessary to discuss an ontology or structured format for representing the research.

A related project, CODECHECK~\citep{nust_codecheck_2021}, aims to provide the verification of the reproducibility of data analysis by a third party in scientific publishing.
CODECHECK proposes a procedure similar to a peer review system, in which the workflow associated with research articles is verified at the time of publication by a reviewer called a CODECHECKER.
However, this project focuses on increasing the availability of the workflow, and does not verify the execution results.
As such, it is unlikely to address the case of our concern that the execution results are not exactly the same, but the conclusions of the study remain the same.
Our proposed metrics, a reproducibility scale of workflow execution results, would be useful in such workflow reproducibility validation in publishing as well.

Software begins to degrade from the moment it is developed, and it is not easy to maintain the same quality over time.
Cases in which an error, including a stack trace, is thrown are quite fortunate; in many cases, the software cannot be executed in the first place, the process does not finish, or the output is inaccurate without throwing an error.
Dealing with such cases and improving debuggability is accomplished by packaging the expected behavior of the software at the time it is developed.
In our approach, we were able to attach information, such as OS, CPU architecture, and dependent software versions, to the expected workflow provenance due to RO-Crate's extensibility.
However, when analysis tools are used internally, as in a workflow, the behavior of the analysis tools tends to be a black box.
Therefore, if an option to display the reproducibility of the execution for each analysis tool is provided, it will be possible to identify the cause of the irreproducibility of the workflow execution results.

We used the file extension to determine the file type and specific tools to extract the biological interpretation from the workflow output.
However, using the file extension is not always reliable, and extracting the biological interpretation from the file contents is not always possible.
Thus, we also tried to take advantage of the summaries and logs generated during the execution of the analysis tools; however, we abandoned this approach due to the diversity and unstructured nature of the summaries and logs.
MultiQC is an existing project that attempts to summarize the results of multiple analysis tools~\citep{ewels_multiqc_2016}.
We believe that integrating our approach with MultiQC will allow us to extract richer information from the workflow output and improve verification accuracy.

In the article regarding a ``reproducibility crisis,'' Baker quoted a Johns Hopkins microbiologist as stating, \textit{``The next step may be identifying what is the problem and to get a consensus.''}~\citep[p.~452]{baker_1500_2016}.
Subsequently, the proliferation of virtualization technology and workflow systems has lowered the bar for re-executing data analysis that an individual or others have previously built.
Despite this, workflow developers are always anxious about whether their workflows are broken.
In response to this anxiety, we realized that the cause is our binary view of whether the workflow could be reproduced or not.
To remove this anxiety, we proposed a new approach to verify the reproducibility of workflows by providing a range of reproducibility of execution results.
With the development of sharing platforms, workflow sharing has become more active.
Therefore, we hope that by verifying reproducibility and sharing the results, more workflows will be reused with confidence, which, in turn, will lead to increased scientific progress.

\section*{Availability of source code and requirements}

\begin{itemize}
    \setlength{\itemsep}{0.2mm}
    \item Project name: Tonkaz
    \item Project home page: \url{https://github.com/sapporo-wes/tonkaz}
    \item DOI: 10.5281/zenodo.7102376
    \item biotoolsID: tonkaz
    \item Operating system(s): Platform independent
    \item Programming language: TypeScript
    \item Other requirements: Deno
    \item License: Apache License, Version 2.0
\end{itemize}

\vspace{2mm}

\begin{itemize}
    \setlength{\itemsep}{0.2mm}
    \item Project name: Sapporo-service
    \item Project home page: \url{https://github.com/sapporo-wes/sapporo-service}
    \item DOI: 10.5281/zenodo.7088999
    \item biotoolsID: sapporo-service
    \item Operating system(s): Platform independent
    \item Programming language: Python
    \item Other requirements: Docker recommended
    \item License: Apache License, Version 2.0
\end{itemize}

\vspace{2mm}

\begin{itemize}
    \setlength{\itemsep}{0.2mm}
    \item Project name: Yevis-cli
    \item Project home page: \url{https://github.com/sapporo-wes/yevis-cli}
    \item DOI: 10.5281/zenodo.7088957
    \item biotoolsID: yevis-cli
    \item Operating system(s): Platform independent
    \item Programming language: Rust
    \item Other requirements: Docker recommended
    \item License: Apache License, Version 2.0
\end{itemize}

\section*{Availability of supporting data and materials}

The workflow, result, and documentation related to the experiment described in the Section ``Results'' are available on GitHub and Zenodo as follows:

\begin{itemize}
    \setlength{\itemsep}{0.2mm}
    \item Execution method and result~\citep{suetake_tonkaz-test-methods_2022}
    \item Workflow definitions~\citep{suetake_test-workflow_2022}
    \item Raw data of workflow execution results~\citep{suetake_tonkaz-raw-data_2022}
\end{itemize}

\section*{Declarations}

\subsection*{List of abbreviations}

ACM: Association for Computing Machinery;
CPU: Central Processing Unit;
CWL: Common Workflow Language;
GA4GH: Global Alliance for Genomics and Health;
ID: Identifier;
OS: Operating System;
RO-Crate: Research Object Crate;
SAM: Sequence Alignment/Map;
VCF: Variant Call Format;
WDL: Workflow Description Language;
WES: Workflow Execution Service;

\subsection*{Ethical Approval}

Not applicable for this study.

\subsection*{Consent for publication}

Not applicable for this study.

\subsection*{Competing Interests}

The authors declare that they have no competing interests.

\subsection*{Funding}

This study was supported by JSPS KAKENHI (Grant Number 20J22439, assigned to H.S.), the CREST program of the Japan Science and Technology Agency (Grant Number JPMJCR17A1, assigned to T.I.), and the Life Science Database Integration Project, NBDC of Japan Science and Technology Agency.

\subsection*{Author's Contributions}

H.S. and T.O. conceived and developed the methodology and software and conducted the investigation.
H.S., T.F., and T.O. wrote the manuscript.
T.F., T.O., and T.I. supervised the project.
All authors read and approved the final version of the manuscript.

\section*{Acknowledgements}

We acknowledge and thank the following scientific communities and their collaborative events where several of the authors engaged in irreplaceable discussions and development throughout the project: the Pitagora Meetup, Workflow Meetup Japan, NBDC/DBCLS BioHackathon Series, and RO-Crate community.
We also acknowledge Prof. Masahiro Kasahara for his valuable comments on the project.

\section*{References}

\bibliography{main}

\end{document}
