## Supplementary figures and images for "A workflow reproducibility scale for automatic validation of biological interpretation results"

### tonkaz_overview.pdf

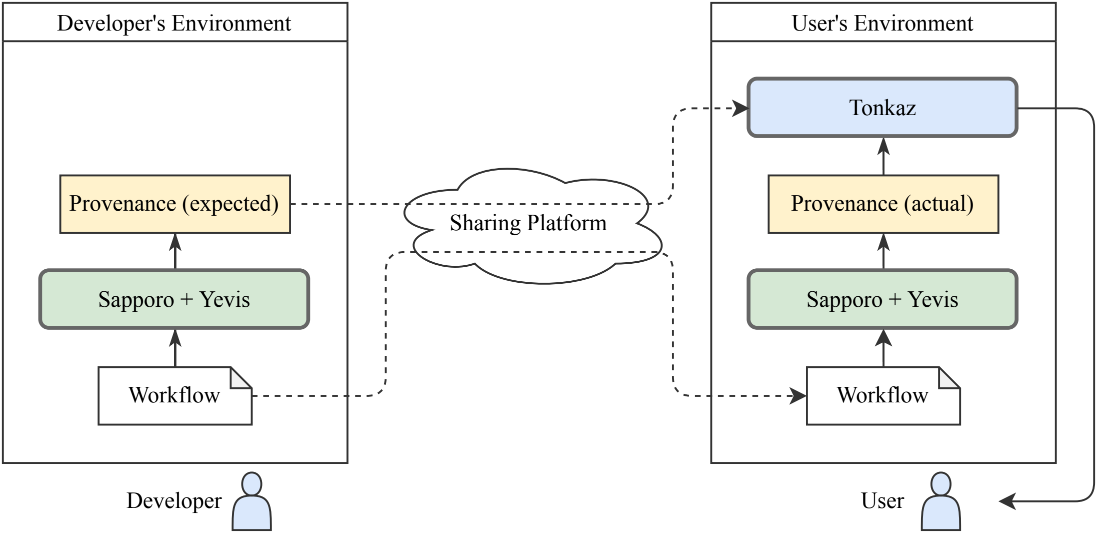

### tonkaz_repro_scale.pdf

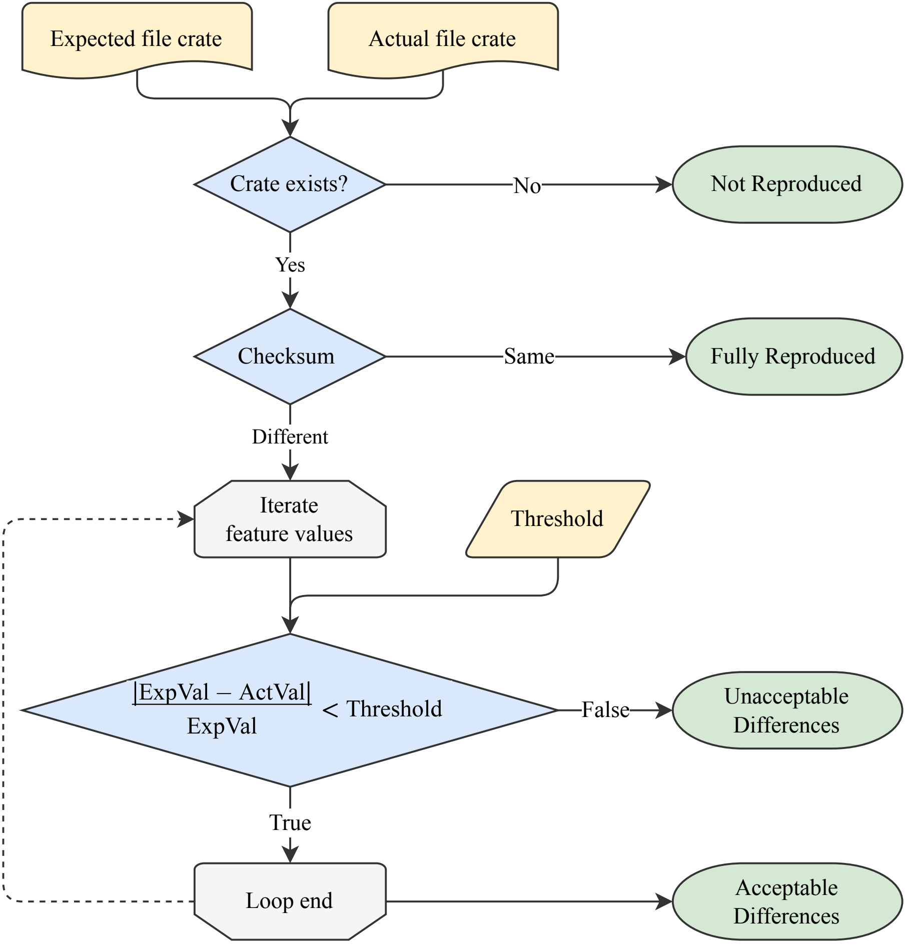
